# The G57 genotype of the BJ/94-like H9N2 lineage exhibits increased replication and virulence in chickens compared to the G1 Middle East Group B lineage

**DOI:** 10.1101/2025.05.28.656345

**Authors:** Sushant Bhat, Jean-Remy Sadeyen, Jiayun Yang, Klaudia Chrazstek, Thusitha K Karunarathna, Mehnaz Qureshi, Dagmara Bialy, Holly Shelton, Munir Iqbal

## Abstract

Avian influenza H9N2 viruses cause significant economic losses to the poultry industry and pose a public health risk due to their potential to reassort with other avian influenza viruses, generating strains with zoonotic and pandemic potential. Two major H9N2 lineages dominate globally: the G1 lineage (genotype G1-B), prevalent in the Middle East, Africa and the Indian subcontinent, and the BJ/94 lineage (predominantly genotype G57), dominant in China, Vietnam, South Korea, Indonesia, and the Far East. We investigated replication, transmission, and pathogenicity of representatives of these two lineages, linking genotype to phenotype. The G57 strain A/Ck/Vietnam/H7F-14-BN4-315/2014 (Vietnam/315) was more lethal to chicken embryos than the G1-B strain A/chicken/Pakistan/UDL-01/2008 (Pakistan/UDL-01). Vietnam/315 exhibited higher replication in both directly infected and contact chickens, with increased virus shedding from the oropharynx and cloaca. In contrast, Pakistan/UDL-01 virus was primarily shed from the oropharynx, highlighting differences in replication, tissue tropism and transmission. Gene analysis showed the M gene of Vietnam/315 enhanced replication in primary chicken kidney cells, whereas the PB2, HA, NA, and M genes promoted increased replication in Madin-Darby Canine Kidney cells. Both viruses showed preferential binding to avian-like receptors over human-like receptors. However, Vietnam/315, however, exhibited higher neuraminidase activity and a more acid-stable HA (pH fusion 5.2) than Pakistan/UDL-01 virus. These findings suggest G57 genotype viruses possess greater replication and transmission fitness than G1-B viruses *in vivo*, ex vivo and *in vitro*. Reassortment events involving G1-B strains acquiring G57 genes may enhance replication and virulence, potentially increasing the risk of animal and human infection.

**Importance:** H9N2 avian influenza viruses are widespread in poultry, resulting in significant economic losses and occasional human infections. Different genetic variants dominate in various regions, but their ability to cause disease and spread in poultry remains unclear. These viruses can exchange genes with other avian influenza strains, altering their infectivity and transmission. This study compared two major H9N2 genotypes: G1-B (common in the Indian subcontinent and the Middle East) and G57 (dominant in China and Vietnam). The G57 virus showed higher replication in laboratory tests and infected chickens, shedding more virus through the orofecal route. It also exhibited a stronger attachment to bird cells and had a more acid-stable HA, suggesting an increased potential for infection and spread.

Our findings indicate that G57 genotype viruses are more infectious and adaptable in poultry than G1 viruses. Gene exchange with other avian influenza strains may generate more virulent viruses with increased transmission potential. This study supports the risk assessment of emerging strains and enhances disease mitigation strategies.

## Introduction

The H9N2 subtype of avian influenza virus (AIV) was first reported in 1966 from turkey farms in the United States [1]. The virus was later identified in poultry in China and diversified into different lineages, which are now prevalent across wider geographical regions worldwide [2]. Although classified as a low pathogenicity AIV, H9N2 outbreaks cause substantial economic losses to the poultry industry through both lethal co-infections and reduced egg production. Additionally, more than 137 zoonotic infections with H9N2 viruses have been reported in many countries, including China, Egypt, Bangladesh, Cambodia, Oman, Pakistan, India, Senegal, and Vietnam [3]. These viruses also show a high tendency to exchange their gene segments with other subtypes of AIVs through reassortment, leading to the emergence of reassortant viruses, often with an expanded host range [4].

H9 haemagglutinin is classified broadly into two phylogeographic lineages: Eurasian and American, with the latter rarely reported [2]. The Eurasian lineage is further divided into three sublineages: G1 (Eastern and Western, found in the Indian subcontinent, Middle East, and Africa) [5,6], BJ/94 (Y280 and G9-like viruses, dominant in China, Vietnam, South Korea and Indonesia) [7], and Y439 (Korean lineage, previously detected in Korea and parts of Eurasia but not reported in recent years) [8].

Both G1 and BJ/94-like viruses have caused human infections [9–11], and extensive genetic reassortment has driven their evolution [12,13]. However, BJ/94-like viruses show higher genetic compatibility for reassortment than G1-like viruses. The G57 genotype is the fittest and the most predominant genotype of H9N2 viruses in China [14,15]. It contributed to the emergence of the highly zoonotic H7N9 virus by donating six internal gene segments [16]. Additionally, at least twelve other subtypes, including H3N8, H5N6, and H10N3, that are linked to human infections acquired one or more internal genes from BJ/94-like H9N2 viruses [17].

Poultry outbreaks with G57 genotype-like viruses have also been reported in other neighbouring countries, including Korea, Myanmar, Tajikistan, and Japan [17]. While G1-like viruses are known to co-circulate with G57 viruses in China and Vietnam, it remains unclear whether G1 viruses are present in all these countries where G57 viruses are enzootic. The G1-like viruses in the Indian subcontinent have evolved by acquiring point mutations that modulate antigenicity [18]. Since their identification in 2008, when G1 viruses were found to have reassorted between HPAI H7N3 and H9N2, resulting in a novel genotype G1-B [5,13], their genotypic constellations have exhibited limited divergence. It is therefore predicted that further genetic reassortment between the co-circulating genotypes, G1-B and G57, could give rise to new variants with greater fitness.

This study assessed the phenotypic characteristics of the BJ/94-like G57 H9N2 virus, which has caused severe disease outbreaks in Vietnam since 2014. The H9N2 strain A/chicken/Vietnam/H7F-14-BN4-315/2014, isolated from a live bird market in northern Vietnam [19], was selected as a prototype virus. Its replication competence in cultured cells, infectivity and virulence in chicken embryos, as well as pathogenicity and transmission potential in chickens, were analysed and compared to the representative G1-B genotype of H9N2 virus A/chicken/Pakistan/UDL-01/2008, isolated from poultry outbreaks in Pakistan in 2008 [5]. Key viral fitness determinants, including pH fusion, receptor binding specificity, and neuraminidase activity, were examined. To identify gene segments responsible for the differing phenotypic characteristics of G57 and G1-B viruses, reassortant viruses were generated using reverse genetics, and their replication phenotypes were analysed using *in vitro* and *ex vivo* replication assays.

## Materials and methods

### Ethics Statement

All studies involving animals (*in ovo* and *in vivo*) were conducted in strict accordance with United Kingdom Home Office regulations and the Animals (Scientific Procedures) Act 1986 under project license number P68D44CF4. All animal work was reviewed and approved by the Animal Welfare and Ethical Review Body at The Pirbright Institute.

### Viruses

The nucleotide sequences for the isolate A/chicken/Pakistan/UDL-01/2008 (hereafter referred to as Pakistan/UDL-01 virus) were retrieved from the National Center for Biotechnology Information (NCBI) (https://www.ncbi.nlm.nih.gov/) (Accession numbers CY038455 to CY038462). The sequences for virus strain A/chicken/Vietnam/H7F-14-BN4-315/2014 (hereafter referred to as Vietnam/315 virus) were retrieved from the National Center for Biotechnology Information (NCBI) (https://www.ncbi.nlm.nih.gov/) (Accession number MH560250.1, MH560222.1, MH560194.1, MH560170.1, MH560143.1, MH560114.1, MH560087.1, MH560068.1). The virus gene segments were synthesised by Geneart^TM^ (Thermo-Fisher Scientific) and subcloned into the pHW2000 vector by restriction enzyme-dependent [20] or restriction enzyme-independent cloning [21,22]. The wild-type viruses and the 1+7 single gene reassortants were rescued by reverse genetics as described previously [23,24] and propagated in 10-day-old specific pathogen-free (SPF) embryonated chicken eggs (ECEs) at 37 °C for 72 hours. The viruses were aliquoted and stored at -80°C until further use.

### Cell lines

The Madin-Darby canine kidney (MDCK) and human embryonic kidney (HEK) 293T cells (ATCC) were maintained with Dulbecco’s Modified Eagle’s medium (DMEM) (Sigma), supplemented with 10% fetal calf sera (FCS) (Sigma), 100 U/mL penicillin-streptomycin (P/S), and 100 μg/mL streptomycin (Gibco). Primary chicken kidney (CK) cells were prepared by the Central Service Unit (CSU) at the Pirbright Institute as previously described [25]. The CKCS were cultured with Eagle’s Minimum Essential Medium (EMEM) (Sigma), supplemented with 2 mm L-glutamine, 0.6% bovine serum albumin (BSA), 1% P/S and 10% tryptose phosphate broth (TPB). All cells were maintained at 37°C under 5% CO_2_.

### Estimation of 50% Embryo infectious dose (EID_50_), Embryo lethal dose (ELD_50_) and replication titers in eggs

The infectious and lethal doses of the viruses to 10-day-old chicken embryos *in ovo* were calculated by making tenfold serial dilutions of the viruses in phosphate-buffered saline (PBS). Each dilution of the virus (100 µL dose) was inoculated into a group of six 10-day-old embryonated hen’s eggs and incubated in an egg incubator maintained at 37°C with 45% humidity for 72 hours. Embryo viability and humane endpoint for culling were determined by candling the eggs twice daily. Embryos showing loss or severe reduction of motility, signs of blood vessel degeneration, and/or detachment of the chorioallantoic membrane (CAM) were considered the humane endpoint and eggs were chilled at 4 °C. Haemagglutination tests prescribed by WOAH [26] were used to determine the viral replication titers of the allantoic fluid harvested from inoculated eggs. Allantoic fluid that was positive for haemagglutination was marked as ‘+’, while allantoic fluid that was negative for haemagglutination was marked as ‘-‘. For the calculation of embryo lethal dose, all embryos that died or reached human endpoints and were positive for haemagglutination were marked as ‘+’. The embryos that showed no signs of severe infection or mortality were marked as ‘-‘. The virus EID_50_ and ELD_50_ were calculated using the method described by Reed and Muench [27].

### In vivo viruses tissue dissemination and transmission from infected to contact chicken

Three-week-old Specific Pathogen-Free (SPF) chickens of the Bovans Brown variety (Henry Stewart & Co., Norfolk, UK) were used in the experiment. For the virus infections, two groups of ten chickens were infected intranasally (i.n.) with 100 µL of inoculum containing 1×10^6^ pfu/mL of either Pakistan/UDL-01 H9N2 or Vietnam/315 H9N2 virus. To study virus transmission to contact chickens, naïve chickens (n = 10) were introduced for co-housing with directly infected chickens at one day post-infection (dpi) and were referred to as ‘early contact chickens’. To identify viral dissemination in the internal organs, directly infected chickens (n = 5) from either group were culled at four dpi. To determine if the virus shed in the later days was transmissible, second-contact chickens (n=4) were co-housed with the directly infected and contact chickens at four dpi and are referred to as ‘late contacts’. The oropharyngeal and cloacal swabs were collected daily in Virus Transport Medium (VTM) from all directly infected chickens, early-contact chickens, and late-contact chickens until eight dpi. A fourth group of six chickens was left uninfected and served as the negative control. Chickens were monitored twice a day to observe the clinical signs. All the chickens were finally swabbed and humanely euthanised by an overdose of pentobarbitone at 14 dpi, and blood was collected adopting terminal cardiac bleed procedure.

### Replication kinetics of H9N2 viruses in cultured cells

Replication kinetics of the wild-type H9N2 viruses and their single gene reassortants were assessed in primary chicken kidney cells (CKCs) and MDCK cells in quadruplicate, using 12-well tissue culture plates [23,28]. For MDCK infection, cells were infected with 0.0001 multiplicity of infection (MOI) of the respective viruses. For CKC infection, cells were infected with 0.0002 MOI in DMEM containing antibiotics and 0.3% Bovine Serum Albumin (BSA). For MDCK cells, the infection medium was supplemented with N-tosyl-L-phenylalanine chloromethyl ketone (TPCK) treated trypsin (2 µg/mL), and the cell supernatant was harvested at 16, 24, 48, and 72 hours post-infection. For CKC cells, the infection medium did not contain TPCK-trypsin, and the cell supernatant was harvested at 19 hours (as it was not feasible to collect at 16 hours as done for MDCKs), followed by 24, 48, and 72 hours post-infection. Viral titers in these harvested samples were determined using standard plaque assay in MDCK cells [29].

### Polymerase activity of H9N2 viruses by minireplicon assay

The polymerase activity of the viral ribonucleoprotein complex in various minigenome combinations of Pakistan/UDL-01 and Vietnam/315 H9N2 viruses was assessed using an *in vitro* minireplicon assay, as described previously [30]. Briefly, the polymerase activity of Pakistan/UDL-01 and Vietnam/315 viruses were compared. Mutations in the PB2 gene unique to the Vietnam/315 virus were introduced into the Pakistan/UDL-01 virus’s PB2 via site-directed mutagenesis to match the corresponding amino acids in the Vietnam/315 virus’s PB2. Each PB2 mutant pCAGGS plasmid (80 ng) was co-transfected into HEK-293T cells alongside ribonucleoprotein expression plasmids for PB1 (80 ng), PA (40 ng), and NP (160 ng) using Lipofectamine 2000 (Invitrogen). Additionally, a pCAGGS plasmid (40 ng) encoding Renilla luciferase and a pCk-PolI-Firefly reporter plasmid (80 ng) encoding firefly luciferase were included in the transfection. Transfected cells were incubated at 37°C for 24 hours, and luciferase activity was measured using the Dual-Glo Luciferase Assay System (Promega). Relative polymerase activity was normalized to that of Renilla luciferase.

### Serology

Seroconversion in the infected chickens was determined via the WOAH-prescribed standard Haemagglutination Inhibition (HI) test [26] using four hemagglutination units of Pakistan/UDL-01 or Vietnam/315 viruses. The reciprocal of the highest dilution of serum showing inhibition of agglutination by chicken red blood cells was taken as the HI titer.

### Plaque morphology

The plaque morphology of the parental viruses and the 1+7 single-gene reassortant viruses was assessed after standard plaque assay [29] using agar overlay on infected MDCK cells. The infected cells were incubated for 72 hrs for plaques to appear and grow. Randomly selected countable plaques (between 9 and 52) were evaluated for their morphology, and plaque size was calculated using ImageJ software [31].

### pH stability of viruses

The acid stability of the H9N2 viruses was assessed by syncytia formation assay in Vero cells under varying pH conditions [32]. Briefly, Vero cells were infected at two-fold dilutions, and the highest virus dilution that infects 100% of cells was determined via immunostaining and used further in the assay. Vero cells pre-seeded in a 96-well plate were infected with H9N2 viruses and incubated for 16 hrs. The infected cells were treated with TPCK-trypsin (3 μg/ml) for 15 minutes and then exposed to PBS buffers (pH 5.2–6.0) for 5 minutes. The buffers were replaced with DMEM (containing 10% FCS), and the plate was incubated at 37°C for 3 hours. The cells were then fixed with ice-cold methanol-acetone (1:1) for 12 minutes and stained with 20% Giemsa stain at room temperature for 1 hour. The pH of fusion was estimated by identifying the lowest pH that induced visible syncytium formation [33]. Images were captured on the Evos XL cell imaging system (Life Technologies).

### Receptor binding

Virus purification and biolayer interferometry were carried out following previously established protocols [18]. Briefly, the H9N2 IAVs were ultracentrifuged at 135,200□×□g for 2 hours and subsequently purified using a continuous sucrose gradient ranging from 30% to 60%. The purified viruses were quantified by indirect ELISA using anti-NP monoclonal antibodies and assessed for receptor binding using an Octet RED bio-layer interferometer (Pall ForteBio, California, USA). The virus was diluted in an HBS-EP buffer (Teknova) supplemented with 10 μM oseltamivir carboxylate (Roche) and 10 μM zanamivir (GSK) to achieve a final concentration of 100 pM. The analysis involved sialoglycopolymers, including α2,6-sialyllactosamine (6SLN), α2,3-sialyllactosamine (3SLN), and Neu5Ac α2,3Gal β1-4(6-HSO3)GlcNAc [3SLN(6-su)], as outlined in previous studies [33]. The relative dissociation constant (Kd), as a measure of binding to 6SLN, 3SLN, and 3SLN(6Su), was calculated.

### Neuraminidase activity

Neuraminidase (NA) inhibition assays were performed on viruses using the fluorogenic substrate 2′-(4-Methylumbelliferyl)-α-D-N-acetylneuraminic acid (MUNANA), as described in a WHO-recommended protocol [34] with minor modifications. Viruses were normalised to 10^5^ PFU and then serially diluted twofold in MES buffer (32.5 mM MES, 4 mM CaCl2, pH 6.5). The MES buffer-virus mixture was then incubated with 100 μM MUNANA substrate at 37°C for 1 hour. The stop solution (0.1 M glycine, 25% ethanol, pH 10.7) was added, followed by measurement using a VICTOR X plate reader (PerkinElmer) with excitation at 355 nm and emission at 460 nm. The EC_50_ of the viruses was calculated as described previously [35] and plotted using Prism version 8.00 for Windows (GraphPad Software, La Jolla, CA, USA).

## Results

### The G57 genotype (Vietnam/315) H9N2 virus exhibits significantly higher replication in embryonated chicken eggs and greater embryo lethality compared to the G1-B genotype (Pakistan/UDL-01) H9N2 virus

The H9N2 viruses from the G1-B genotype (Pakistan/UDL-01) and G57 genotype (Vietnam/315) used in this study were rescued by reverse genetics and propagated in Embryonated Chicken Eggs (ECEs). The Vietnam/315 virus replicated to significantly higher levels in SPF ECEs than the Pakistan/UDL-01 virus, as determined by haemagglutination (HA) testing of allantoic fluid using 1% chicken RBCs (Figure 1A). Moreover, the Vietnam/315 virus exhibited greater lethality, causing nearly 84% embryo mortality within 72 hours, compared to around 33% for the Pakistan/UDL-01 virus (Figure 1B). These findings indicate that G57 genotype viruses replicate to a higher titer in ECEs and are more virulent than G1-B genotype in chicken embryos.

**Figure 1.**
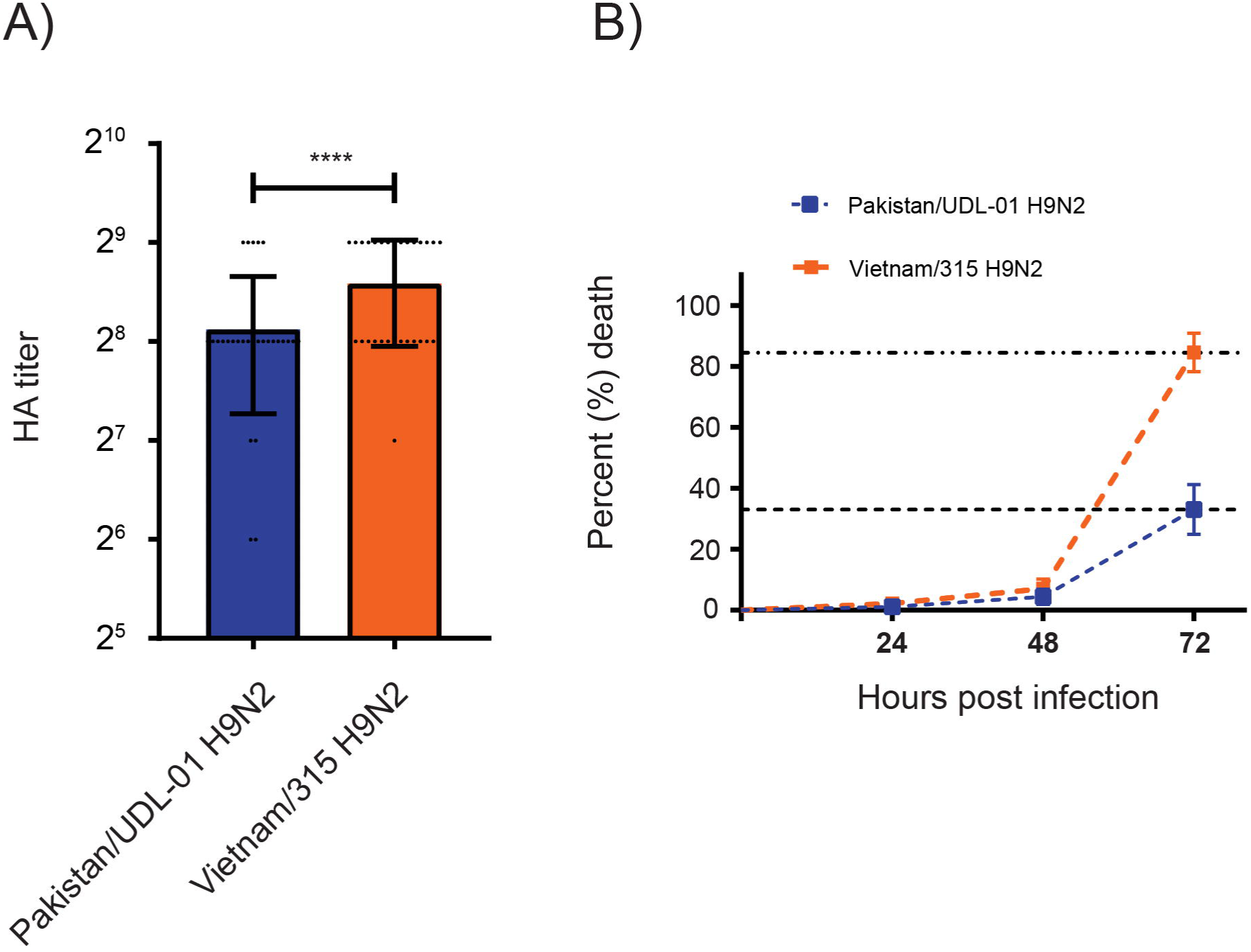
*In ovo* replication and embryo lethality of Vietnam/315 and Pakistan/UDL-01 H9N2 viruses. The replication fitness and lethality of Vietnam/315 and Pakistan/UDL-01 H9N2 viruses were assessed *in ovo* by infecting 10-day-old specific-pathogen-free (SPF) chicken embryos with 200 plaque-forming units (PFU) of the respective virus in a 100 µL dose. (A) Allantoic fluid was harvested 72 hours post-infection and virus replication titers were estimated by haemagglutination test. (B) Virus lethality was assessed by monitoring embryo mortality over 72 hours following infection. The mean viral titers is shown with standard deviation. The two groups were compared using an unpaired t test, with P value = 0.0037 indicated by **

### Vietnam/315 virus exhibits higher viral replication in SPF chickens compared to Pakistan/UDL-01 virus

The productive replication, pathogenesis, and transmissibility of G1 (Pakistan/UDL-01) and G57 (Vietnam/315) H9N2 viruses were compared in 3-week-old SPF chickens. Chickens directly inoculated via the intranasal route (referred to as ‘D0’) with the Pakistan/UDL-01 virus or Vietnam/315 virus exhibited productive infection, shedding the virus primarily through the oropharynx (Figure 2).

**Figure 2.**
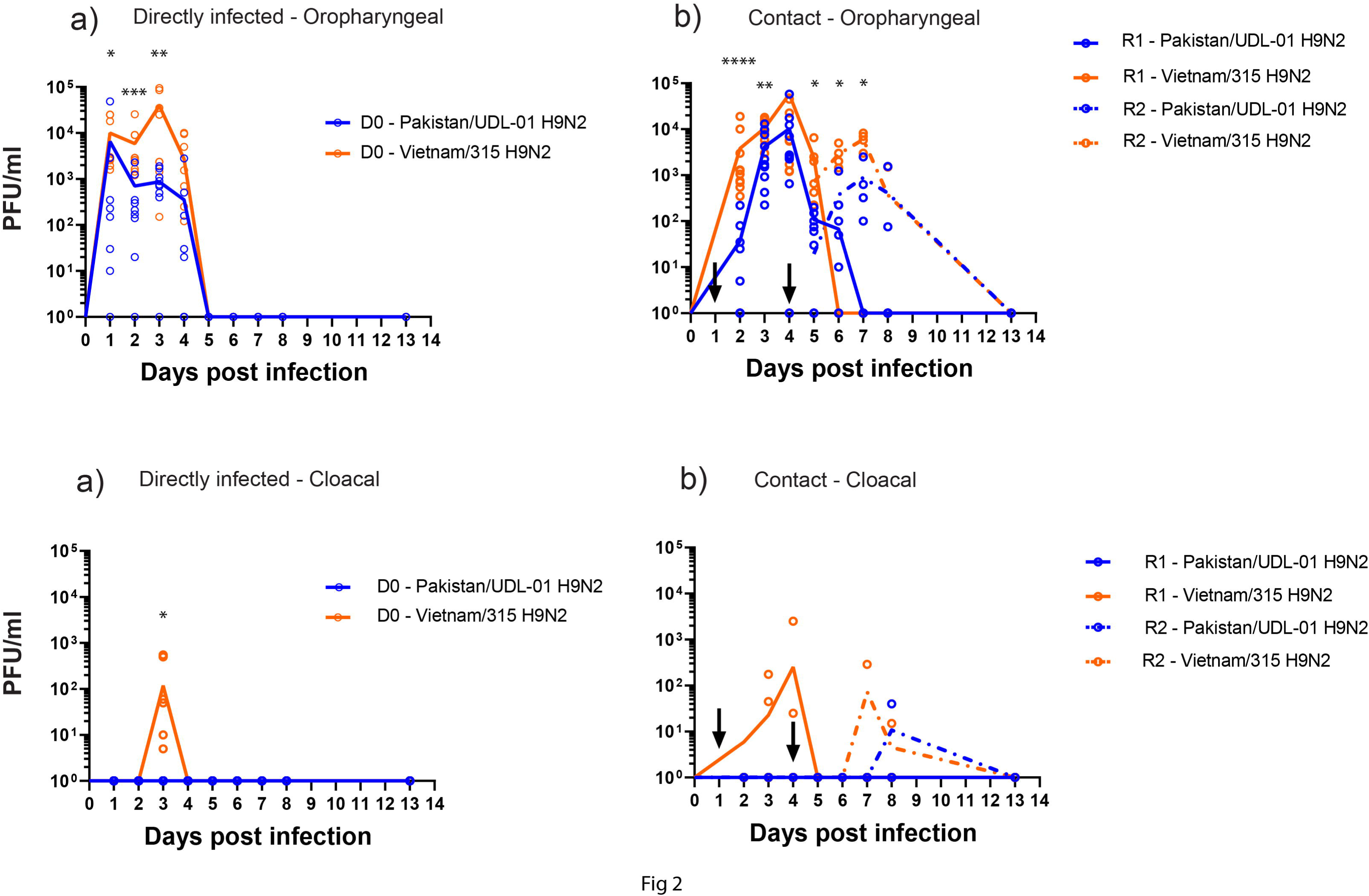
Comparative mean virus shedding levels and transmission dynamics of Pakistan/UDL-01 H9N2 virus and Vietnam/315 H9N2 virus in specific pathogen-free chickens. The virus shedding (shown as PFU/ml) was identified by titrating the swab samples (oropharyngeal and cloacal) taken from chickens directly infected (D0) with the Pakistan/UDL-01 H9N2 or Vietnam/315 H9N2 (a, c) and contact chickens co-housed with the D0 chickens on day 1 post-infection (R1) or day 4 post-infection (R2) (shown as arrows) (b, d). Swab samples were taken daily from the D0, R1 and R2 chickens until day 8 post-infection. All chickens were finally swabbed and humanely sacrificed on day 13 post-infection. Each time point represents the mean viral shedding, with standard deviations indicated. The viral shedding indicated on respective days in the two groups was compared using the Mann-Whitney t test. The asterisks indicate P values; *<0.05, **<0.005, ***<0.0005, ****<0.0001.

The oropharyngeal virus shedding in the ‘D0’ group of chickens infected with Vietnam/315 virus (Figure 2a) exhibited mean peak virus shedding (10^4.59^ PFU/ml) at three dpi, while ‘D0’ group of chickens infected with Pakistan/UDL-01 virus exhibited lower virus shedding with peak viral shedding at one dpi (mean titer – 10^3.81^ PFU/ml).

The first-contact chickens (hereby referred to as ‘R1’) in the Vietnam/315 virus group and the Pakistan/UDL-01 virus group exhibited peak oropharyngeal virus shedding at four dpi (three days post-exposure - dpe) (Figure 2b). However, the peak oropharyngeal virus shedding in the Vietnam/315 virus group was higher (mean titer: 10^4.76^ PFU/ml) than in the Pakistan/UDL-01 virus group (mean titer: 10^3.95^ PFU/ml).

The virus shedding from the oropharynx continued until five dpi (four dpe) and ceased at six dpi (five dpe) in the Vietnam/315 virus group. On the other hand, the virus continued to be shed from the oropharynx until six dpi (five dpe) in the Pakistan/UDL-01 virus group and ceased at seven dpi (six dpe). The second contact chickens (hereby referred to as ‘R2’) in both the groups showed peak oropharyngeal virus shedding at seven dpi (three dpe), with Vietnam/315 virus group showing higher oropharyngeal virus shedding (mean titer – 10^3.78^ PFU/ml) than Pakistan/UDL-01 virus (mean titer – 10^2.95^ PFU/ml). The oropharyngeal virus shedding in the R2 groups continued until eight dpi (four dpe) (Figure 2b), when daily swabbing was stopped.

The cloacal virus shedding in the D0 group was observed only in the Vietnam/315 virus group on three dpi in six out of ten chickens (mean titer – 10^2.07^ PFU/ml), while the D0 chickens in the Pakistan/UDL-01 virus group did not show any virus shedding via cloaca (Figure 2c). The cloacal virus shedding exhibited by R1 contacts in the Vietnam/315 virus group continued until four dpi (three dpe) (mean titer: 10^2.40^ PFU/ml), while the R1 contacts in the Pakistan/UDL-01 virus group did not show any cloacal virus shedding (Figure 2d). The R2 contacts in Vietnam/315 virus group showed peak cloacal virus shedding (mean titer – 10^1.86^ PFU/ml) at seven dpi (three dpe), while R2 contacts in Pakistan/UDL-01 virus group showed peak cloacal virus shedding (mean titer – 10^1.03^ PFU/ml) at eight dpi (four dpe) (Figure 2d). Overall, these results indicate that the Vietnam/315 virus replicates more efficiently, is shed at higher levels in directly infected chickens, and causes more severe infection in sentinel chickens compared to the Pakistan/UDL-01 virus. The Vietnam/315 virus sheds from both the oropharynx and cloaca. In contrast, the Pakistan/UDL-01 virus sheds mainly via the oropharynx, as the mean cloacal shedding in second-contact birds was close to the baseline.

### Vietnam/315 virus shows more dissemination in internal organs in specific pathogen-free chickens compared to Pakistan/UDL-01 virus

To determine the dissemination of H9N2 viruses into internal organs, lungs and kidneys were collected from five directly inoculated chickens per group at four dpi. No virus was detected in chickens infected with Pakistan/UDL-01 virus, whereas four directly inoculated chickens and three contact chickens infected with Vietnam/315 virus showed the presence of the virus in their lungs. Two directly inoculated chickens and two contact chickens showed the presence of the virus in their kidneys (Figure 3). These results indicate that the Vietnam/315 virus caused a more systemic infection compared to the Pakistan/UDL-01 virus.

**Figure 3.**
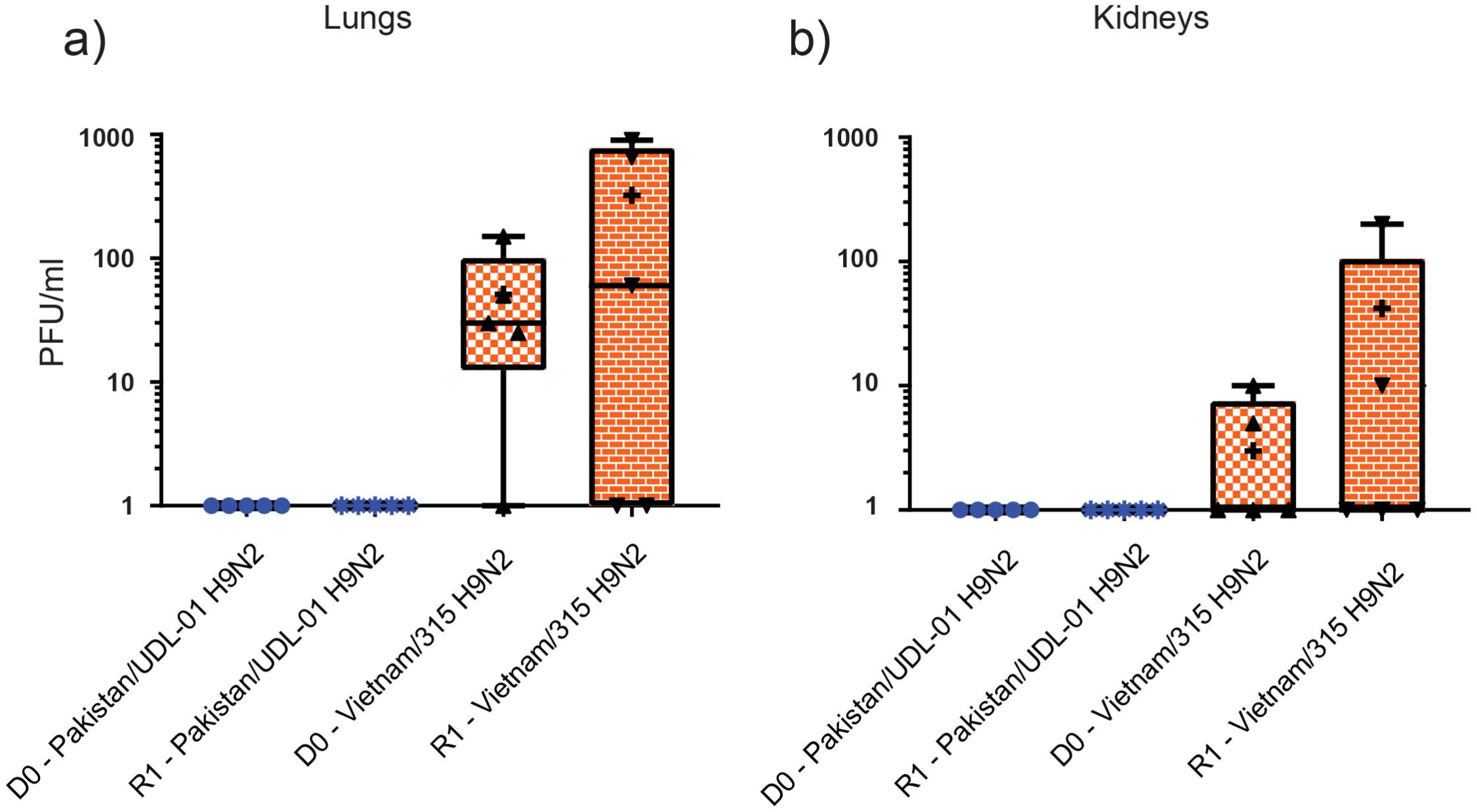
Virus dissemination in lungs (a) and kidney (b) of chickens directly infected with Pakistan/UDL-01 H9N2 and Vietnam/315 H9N2. Four chickens directly infected with the Pakistan/UDL-01 or Vietnam/315 virus were scheduled for post-mortem examination at 4 days post-infection (dpi) to estimate virus levels in tissues. Levels of virus detected in the lungs and kidneys were estimated by plaque assay using 100 mg of tissue. The error bars indicate standard deviation in the viral titers. No statistical significance was identified between directly infected (D0) and contact (R1) chickens.

### The Vietnam/315 virus replicates more efficiently in primary chicken kidney cells compared to the Pakistan/UDL-01 virus, with the difference attributed to the M gene

To determine viral replication *ex vivo*, the H9N2 viruses were compared for their replication in primary Chicken Kidney Cells (CKCs). The Vietnam/315 virus exhibits a significantly higher (P□< 0.0001) replication rate in *ex vivo* cultures of primary Chicken Kidney cells (CKCs) compared to the Pakistan/UDL-01 virus at all time points (19-72 hours) analysed (Figure 4). The Vietnam/315 virus attains a titer around 3×10^6^ pfu at 19 hours post-infection and a peak titer of 8.8×10^7^ pfu at 48 hours post-infection, while the Pakistan/UDL-01 virus attains a titer of 0.63×10^3^ pfu at 19 hours post-infection and a peak titer of 3.5×10^6^ at 48 hours post-infection. The multi-step replication kinetics of single gene reassortants showed that replacing the M gene of Pakistan/UDL-01 virus with Vietnam/315 virus increases the replication at early time points (19 hours and 24 hours post-infection) (P□< 0.0001). However, NS gene replacement reduces the replication at 19 hours (P<0.005), but the difference becomes non-significant as the replication progresses until 72 hours. On the other hand, replacing the PB2, PB1, PA, HA, NP, and NA genes of the Pakistan/UDL-01 virus with those of the Vietnam/315 virus reduces the replication rate in primary CK cells. These results indicate that the increased replication of the Vietnam/315 virus in primary chicken kidney cells is attributed, at least in part, to the M gene.

**Figure 4.**
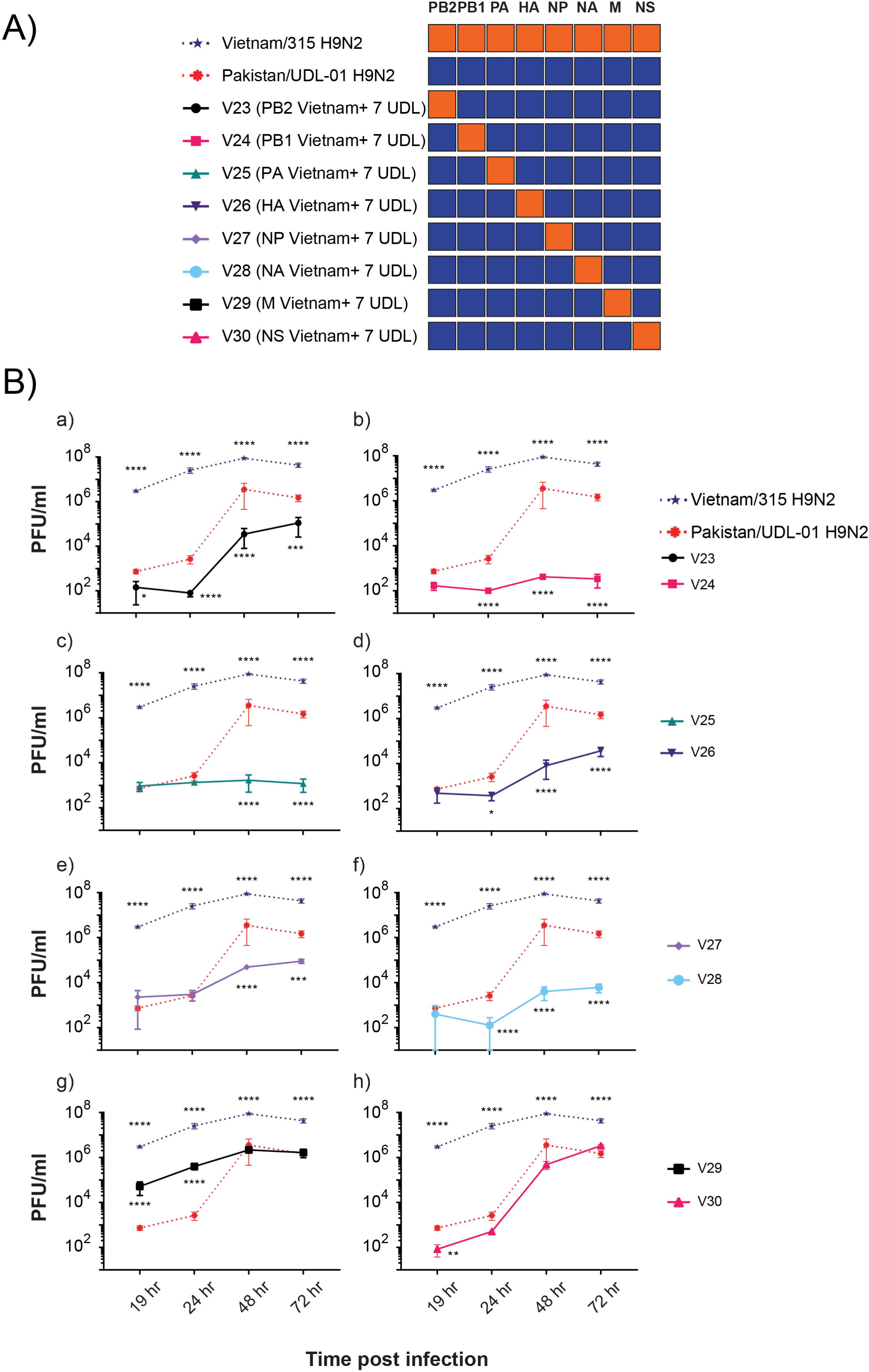
Multistep replication kinetics of parental H9N2 viruses and 1+7 reassortants containing the single gene of Vietnam/315 virus and seven genes from Pakistan/UDL-01 virus in primary Chicken Kidney cells. The contribution of each gene segment of the Vietnam/315 virus to replication fitness was determined by **A)** generating 1+7 reassortant viruses using reverse genetics. Each of the indicated reassortant viruses analysed contained one gene segment from Vietnam/315 virus and the remaining seven gene segments from Pakistan/UDL-01 virus. **B)** The replication of reassortant viruses compared to parental H9N2 viruses is shown in panels ‘a’ to ‘h’. The multistep replication kinetics were determined by infecting primary Chicken Kidney cells with the respective viruses at a 0.0002 multiplicity of infection (MOI). The cell supernatant was harvested at 19, 24, 48, and 72Lhours post-infection. Viral titers were determined by plaque assay using an overlay medium containing 0.7% Avicel. Each time point represents the mean of three biological replicates, with standard deviations indicated. A two-way ANOVA with multiple analyses was performed to compare every group to Pakistan/UDL-01 H9N2. The asterisks: * indicate P values < 0.05 ; ** indicate P values < 0.005 ; *** indicate PL=L0.001; ****, PL< 0.0001.

### The Vietnam/315 virus replicates to a higher titer in MDCK cells and forms relatively larger plaques, which is attributed to the PB2, HA, NA, and M gene segments

The comparative replicative fitness and plaque morphology of Vietnam/315 and Pakistan/UDL-01 Hwere analysed in mammalian-origin cells. The Vietnam/315 virus exhibits more cell-to-cell spread, as indicated by increased plaque size (P < 0.0001) compared to the Pakistan/UDL-01 virus (Figure 5B). The plaque morphology of single-gene reassortants containing the PB2, HA, NA, or M gene of the Vietnam/315 virus with a Pakistan/UDL-01 virus backbone exhibited an increased plaque size in MDCK cells (Figure 5B).

**Figure 5.**
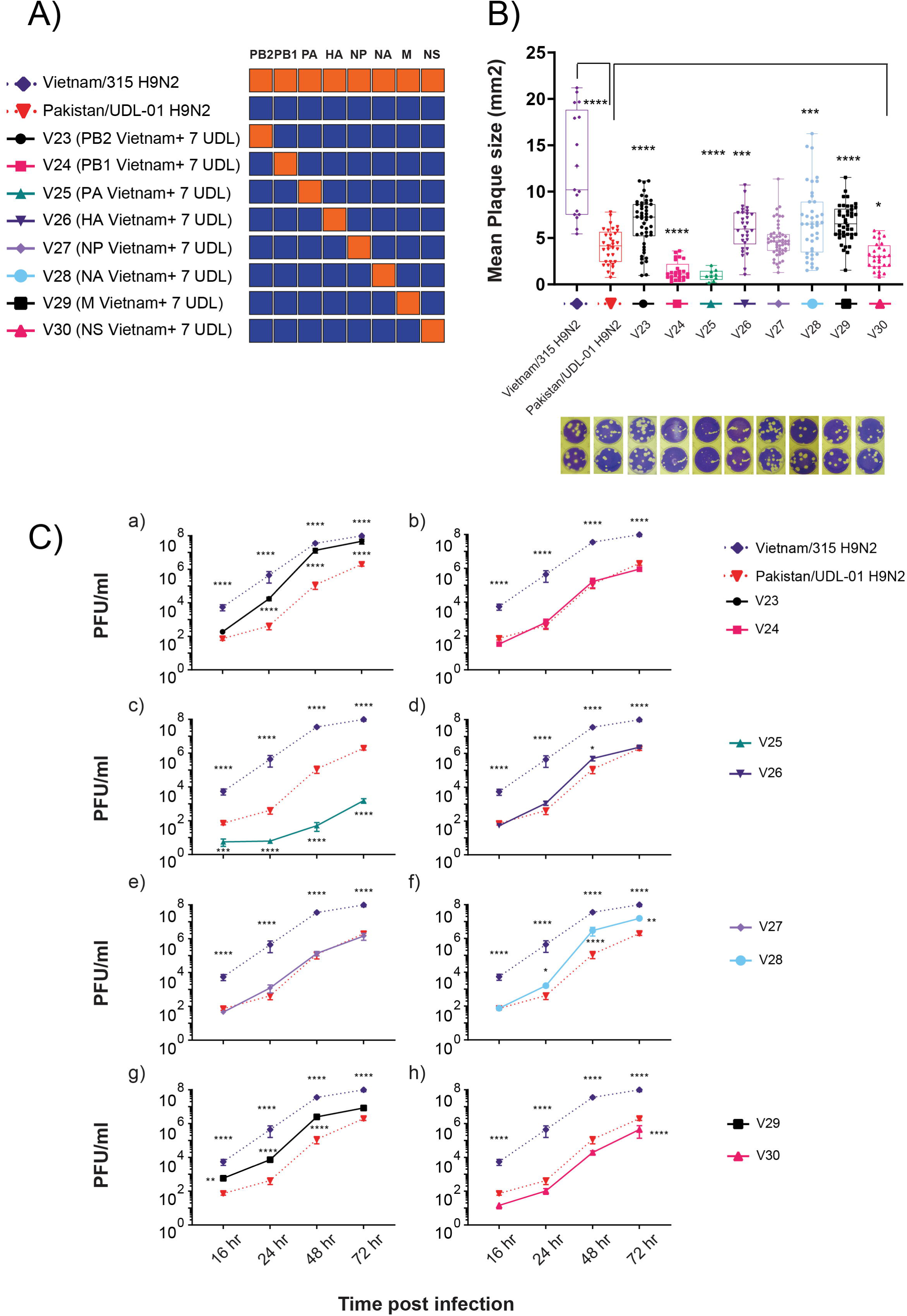
Multistep replication kinetics and plaque morphology of parental H9N2 viruses and 1+7 reassortants containing the single gene of Vietnam/315 virus and seven genes from Pakistan/UDL-01 virus in Madin-Darby canine kidney (MDCK) cells. The contribution of each gene segment of the Vietnam/315 virus to replication fitness was determined by **A)** generating 1+7 reassortant viruses using reverse genetics. Each of the indicated reassortant viruses analysed contained one gene segment from Vietnam/315 virus and the remaining seven gene segments from Pakistan/UDL-01 virus. **B)** Mean plaque size (mm^2^) of the parental H9N2 viruses and the reassortant viruses was calculated in pixels using ImageJ software and converted to mm². The data were plotted along with the Standard Error of Mean (SEM) and compared to the Pakistan/UDL-01 H9N2 virus using Dunnett’s multiple comparison test (one-way ANOVA). The total number of counted plaques ranged from 9 to 52, depending on the number of countable plaques. **C)** The replication of reassortant viruses compared to parental H9N2 viruses is shown in panels ‘a’ to ‘h’. The multistep replication kinetics were determined by infecting MDCK cells with the respective viruses at a 0.0001 multiplicity of infection (MOI). The cell supernatant was harvested at 16, 24, 36, and 48 hours post-infection. Viral titers were determined by plaque assay using overlay media containing 1% Agarose. Each time point represents the mean of four biological replicates, with standard deviations indicated. A two-way ANOVA with multiple analyses was performed to compare every group to Pakistan/UDL-01 H9N2. The asterisks: * indicate P values < 0.05 ; ** indicate P values = 0.0013 ; *** indicate PL=L0.008; ****, PL< 0.0001.

The Vietnam/315 virus showed a significantly higher replication rate (P□< 0.0001) than the Pakistan/UDL-01 virus at all time points in MDCK cells (Figure 5C). The replication kinetics of single-gene reassortants, in which the PB2, HA, NA, or M gene of the Vietnam/315 virus was introduced into the Pakistan/UDL-01 virus backbone, resulted in increased replication in MDCK cells (Figure 5C). This suggests that the enhanced replication and larger plaque size of the Vietnam/315 virus in MDCK cells are attributed to the PB2, HA, NA, and M gene segments.

### The Vietnam/315 virus exhibited more acid-stable HA compared to the Pakistan/UDL-01 virus

The pH fusion of haemagglutinin of H9N2 viruses was estimated by the syncytium formation assay in Vero cells. Consistent with the previous reports [24,36], the Pakistan/UDL-01 virus showed a pH fusion of 5.4. Interestingly, the Vietnam/315 virus showed a lower pH threshold fusing at pH 5.2 or lower. This indicates that the Vietnam/315 virus requires a lower pH for HA to undergo conformational change and trigger endosomal membrane fusion, resulting in a more acid-stable HA compared to the Pakistan/UDL-01 virus.

### Vietnam/315 virus exhibited relatively higher receptor binding avidity towards avian-like sialic acid receptor analogues compared to human-like receptor analogues

The receptor binding specificity of Vietnam/315 virus was quantified and compared with that of Pakistan/UDL-01 virus using synthetic sialoglycopolymers—α2,6-sialyllactosamine (6SLN), α2,3-sialyllactosamine (3SLN), or sulfated 3SLN (6-Su) receptor analogues — by bio-layer interferometry. Binding to 3SLN was not detectable for either virus, indicating negligible interaction. Both strains showed moderate binding to 6SLN, with Pakistan/UDL-01 exhibiting slightly stronger binding (K_d_ = 0.2075 μM) compared to Vietnam/315 (0.3921 μM). In contrast, markedly stronger binding was observed to the modified receptor 3SLN(6Su), with K_d_ values of 0.0046 μM for UDL and 0.0029 μM for Vietnam. This represents a ∼45–135-fold increase in binding affinity relative to 6SLN, highlighting a clear preference for 3SLN(6Su). Although the observed differences between Pakistan/UDL-01 and Vietnam/315 at each receptor are relatively small and may fall within expected assay variability, they suggest subtle differences in receptor-binding dynamics between the two strains. Overall, these data indicate a strong binding preference for 3SLN(6Su) over 6SLN or 3SLN, consistent across both viruses.

### Vietnam/315 virus has a higher enzymatically active NA than Pakistan/UDL-01 H9N2

The contribution of NA to the differential fitness of these H9N2 viruses was assessed by measuring neuraminidase activity using a Fluorometric MUNANA Assay. The Vietnam/315 virus exhibited higher neuraminidase activity, as indicated by lower EC_₅₀_ values, compared to the Pakistan/UDL-01 virus (Figure 6B, a, b). This suggests that a lower concentration of Vietnam/315 virus NA protein can antagonize 50% of the MUNANA substrate, requiring less viral NA for efficient progeny virus release from infected cells.

**Figure 6.**
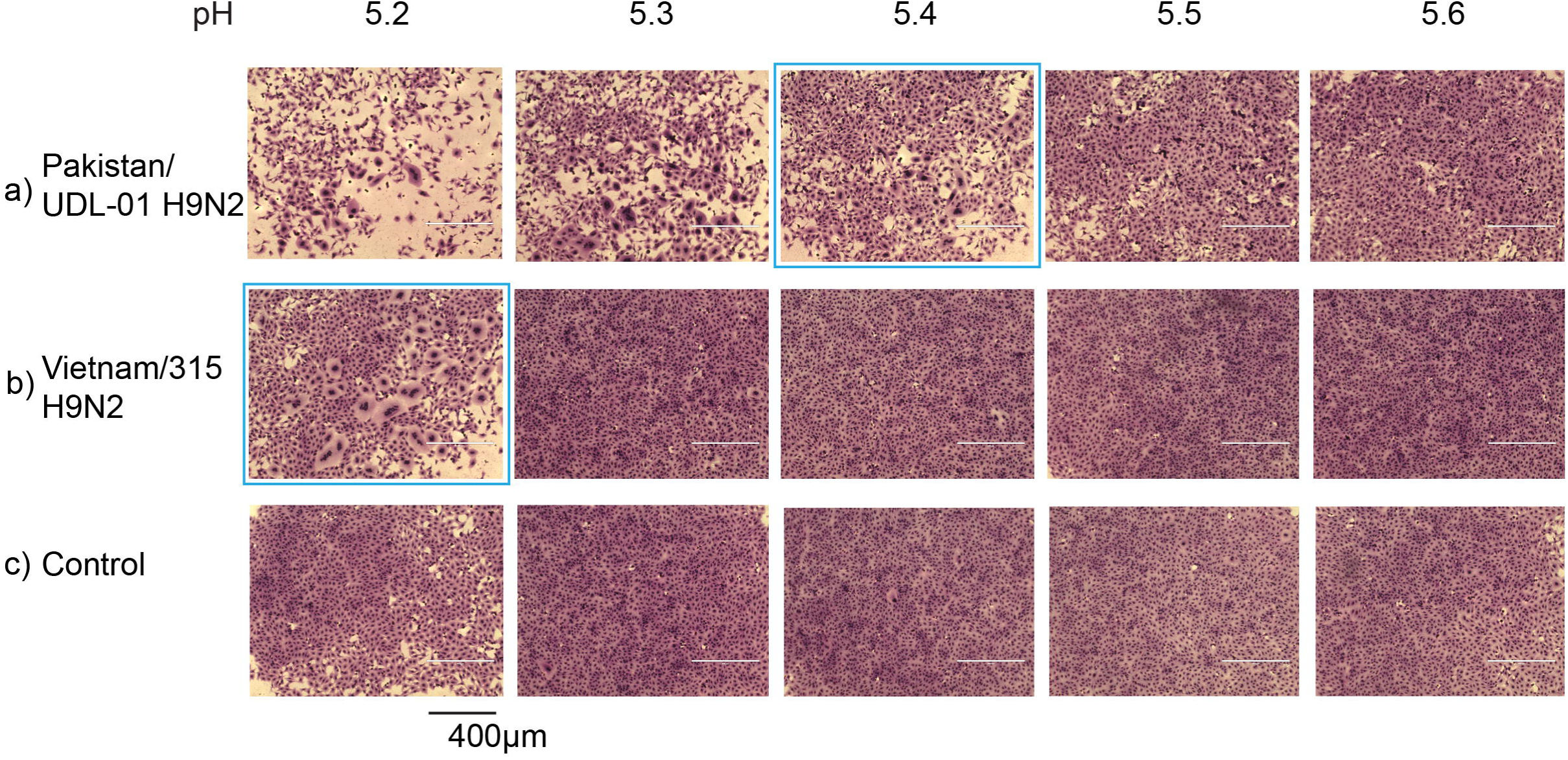
pH fusion of H9N2 viruses estimated by syncytium formation in Vero cells. The cells were infected with a) Pakistan/UDL-01 H9N2 or b) Vietnam/315 H9N2, and syncytium formation was identified by exposing the infected cells to buffers in the pH range of 5.2-6.0 (images shown only up to pH 5.6). The uninfected cells (c) exposed to different pH served as negative control.

**Figure 7.**
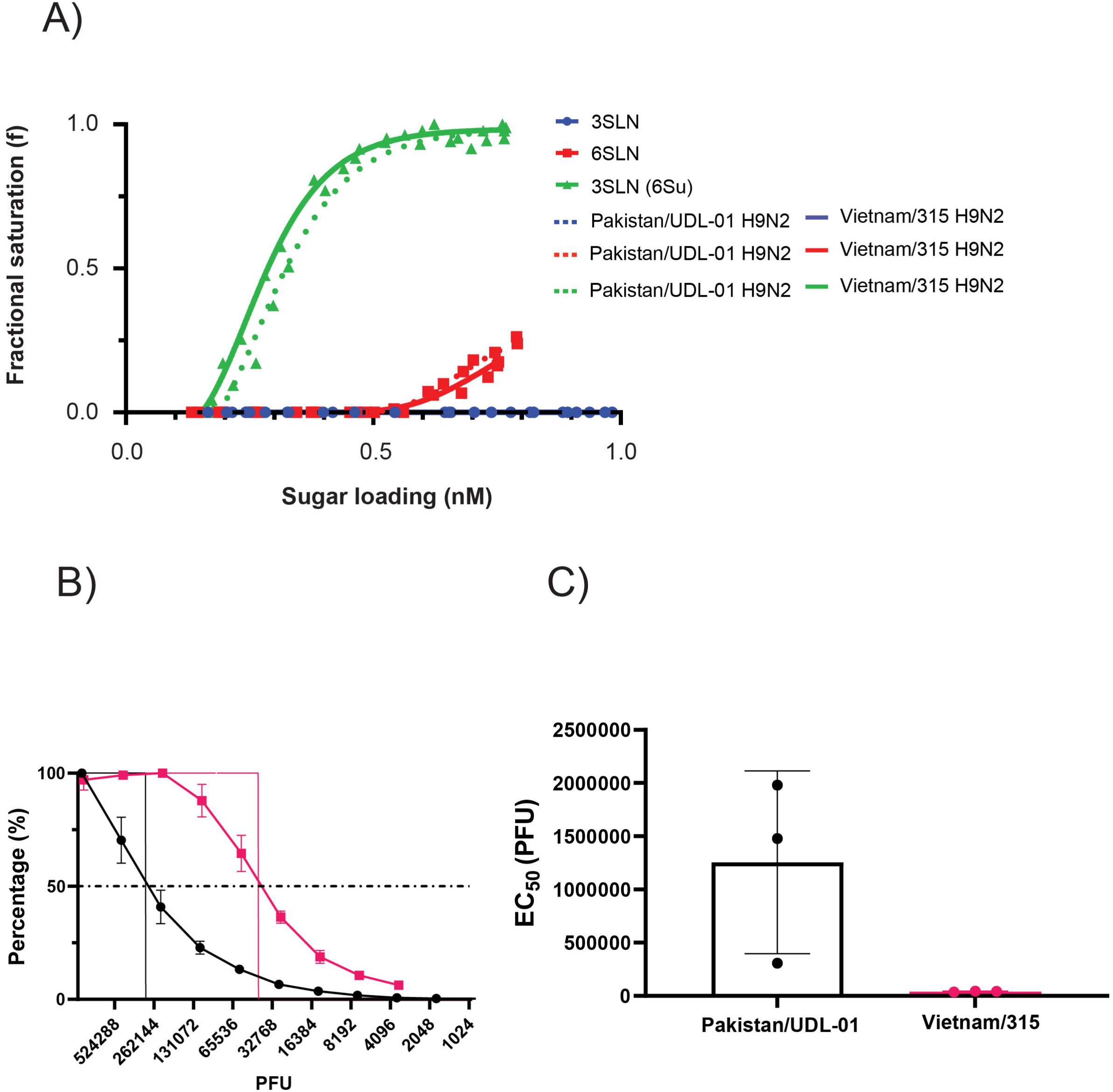
Receptor binding profile and Neuraminidase (NA) sialidase activity of Pakistan/UDL-01 H9N2 and Vietnam/315 H9N2. **A)** The receptor binding profiles of Pakistan/UDL-01 H9N2 and Vietnam/315 H9N2 viruses were determined using biolayer interferometry. The viruses were normalised in terms of NP content, and an equimolar amount of viruses was analysed for receptor binding towards α-2,3-linked (3′SLN 6-sulfated), α-2,3-linked (3′SLN) or α-2,6-linked (6′SLN) sialylglycan receptors **B)** Neuraminidase activity was assessed using MUNANA assay. Viruses were normalized in terms of PFU (10^6^ PFU/ml) and two-fold dilutions of viruses were subjected to MUNANA substrate. The fluorescence activity was plotted against PFU with error bars indicating standard deviation, and EC50 was calculated.

## Discussion

H9N2 avian influenza viruses have undergone significant evolutionary changes since their initial identification [1], leading to alterations in replication efficiency, virulence, transmission dynamics, and zoonotic potential. In the early 1990s, H9N2 strains of both the G1 and BJ/94 lineages primarily caused mild or subclinical infections in chickens [37]. However, due to genetic mutations and reassortments, contemporary genotypes—most notably G1-B and G57— have been associated with more severe disease outbreaks in poultry populations [38,39]. The G1-B genotype of the H9N2 G1 lineage emerged through reassortment with co-circulating highly pathogenic H7N3 viruses in Pakistan. This involved the acquisition of internal gene segments that significantly contributed to its fitness and dominance in poultry across the Indian subcontinent, the Middle East, and Africa, replacing the older G1 lineage viruses [5,13]. Meanwhile, the G57 genotype of the BJ/94 lineage has become prevalent in China, Vietnam, and neighbouring countries, establishing itself as the dominant genotype since its identification in China in 2007 [40]. Although both G1-B and G57 genotypes have maintained dominance in their respective regions, limited data are available on their key phenotypic differences, particularly in replication capacity, tissue tropism, virulence and the molecular factors driving the phenotypic differences between the two genotypes.

We utilised a combination of *in vitro*, *in ovo*, *ex vivo*, and *in vivo* models as well as reverse genetics approaches to compare the replication kinetics, tissue dissemination, and pathogenicity of two representative strains: Pakistan/UDL-01 (G1-B genotype) and Vietnam/315 (G57 genotype). We also identified the molecular factors driving the phenotypic differences between the currently dominant genotypes of H9N2 viruses. The findings demonstrate that the G57 genotype exhibits higher virulence in chicken embryos and a greater dissemination in the lungs and kidneys of infected chickens compared to the G1-B genotype. Molecular analysis revealed that the Vietnam/315 strain possesses a distinct combination of PB2, HA, NA, and M gene segments, which are likely contributors to its increased fitness and pathogenicity.

The genotypic variability of H9N2 strains likely influences the pathogenicity phenotype and tissue dissemination as genetic differences can directly modulate viral receptor binding, neuraminidase activity, polymerase function, and/or the ability to antagonise host antiviral responses [41–43]. *In vitro* studies on the multistep replication kinetics of single-gene reassortants from the Vietnam/315 strain, when combined with the Pakistan/UDL-01 background, revealed that the higher replication of Vietnam/315 virus in avian cells is primarily attributed to the M gene segment. However, it is important to note that epistatic interactions between different gene segments also play a significant role. The reassortment of the M gene contributes to increased viral replication in avian cells, possibly by enhacing the export of ribonucleoprotein from the nucleus [44].

The crucial role of the M gene in viral replication was further demonstrated in our previous study [23], where a reassortant H7N9 bearing PA and M from Vietnam/315 strain was generated by coinfecting chickens with H7N9 and H9N2 and the reassortant virus exhibited increased replication compared to its parental strain.

The Vietnam/315 strain also exhibited a significantly higher replication rate and increased cell-to-cell spread, resulting in larger plaque sizes in MDCK cells when compared to Pakistan/UDL-01. This phenotype can be attributed to its PB2, HA, NA, and M gene segments. Notably, the PB2 and M gene segments in the G57 genotype originated from Y280-like (or BJ/94-like) lineages of H9N2, while the G1 Middle East lineage contains the PB2 and M segments from the H7N3 subtype. Additionally, the PB2 of the Vietnam/315 possesses various amino acid markers associated with higher polymerase activity (Supplementary Fig. S1) and enhanced replication in mammalian cells [45,46] compared to the PB2 of the G1 Middle East lineage.

Receptor binding analyses indicated that both strains preferentially bind sulfated avian 3′SLN receptors. However, Vietnam/315 showed higher neuraminidase activity than Pakistan/UDL-01, suggesting a more efficient viral release contributed by Vietnam/315 NA. This facilitates efficient viral attachment and release from host cells, supported by replication kinetics data showing significantly higher viral titers for Vietnam/315 at 19 hours post-infection. Additionally, our previous work demonstrated that the six internal genes of Vietnam/315 confer a selective advantage to H7N9 viruses, enhancing their replication and transmission in chickens [23]. This advantage is attributed to the higher polymerase activity of the RNP complex, indicating that a combination of an optimized HA-NA balance and a highly active RNP complex confers Vietnam/315 strain a significant phenotypic edge over Pakistan/UDL-01 [14,15,47,48].

While the human seasonal and pandemic influenza viruses often require lower pH (<5.5) for fusion, the avian influenza viruses typically have higher pH fusion [49]. The acid-stable HA of Vietnam/315 can contribute to its stability in respiratory droplets and aerosols, promoting airborne transmission, a characteristic shared with other human-adapted influenza viruses like pdm H1N1 and H3N2. The increased replication and shedding of Vietnam/315 in chickens heightens the risk of spillover to humans. The Vietnam/315 virus also possesses unique mammalian adaptation markers that improve its capacity to interact with human host factors, such as PB2 271I interacting with ANP32 [50] and NP 52N conferring both BTN3A3 evasion [51], and overcoming Mx1 restriction [52]. These markers confer increased susceptibility to replication and immune evasion in the human host, thereby facilitating cross-species transmission and infection. The Vietnam/315 virus, despite demonstrating lower binding avidity for human 6SLN receptors compared to avian receptors, undeniably has the capacity to cause human infections, as evidenced by recent findings with G57 viruses [17]. Any future adaptation enhancing human receptor binding, while retaining the acid stability of HA, could further increase the risk of human transmission.

Genetic characterisation of G57 viruses from Vietnam revealed an HA cleavage site with the dibasic RSSR motif and residues associated with human receptor binding [19]. The Vietnam/315 strain carries S316 (H9 numbering) in HA and a three amino acid deletion in the NA stalk, which are linked to improved HA cleavage and increased virulence in chickens and mice [53]. These features have become increasingly common among H9N2 viruses isolated since 2002 [54] (Supplementary Fig. S2) and contribute to productive infection, enhanced dissemination, and greater viral shedding — ultimately facilitating efficient transmission within poultry populations.

Virus shedding patterns differed between the strains: Vietnam/315 was shed from both the oropharynx and cloaca, with respiratory shedding predominating as is common with H9N2 viruses in chickens [55], while the Pakistan/UDL-01 virus was primarily shed through the oropharynx. While cloacal shedding is more common in waterfowl, it has been occasionally observed in terrestrial poultry infected with certain low pathogenic H9N2 viruses [56]. We have previously observed cloacal shedding in Rhode Island Red chickens infected with Pakistan/UDL-01 virus [57]. It was not significant in the Bovans Brown breed of chickens in the current study, possibly due to breed-dependent susceptibility [58]. Furthermore, low pathogenic viruses with more than dibasic HA cleavage motifs can lead to systemic infections, increasing the likelihood of cloacal shedding and facilitating faecal-oral transmission, as observed with Vietnam/315 [56].

Due to the migratory-wild-domestic bird interface across the major wetlands of the Indian sub-continent [59], reassortant viruses between G1 and G57 lineages have been generated which have an increased propensity to replicate in terrestrial poultry [60]. Our study elucidates the molecular factors responsible for the increased replication and fitness of G57 viruses. This contributes to the risk assessment of emerging reassortant viruses between G1 and G7 genotypes. Therefore, continuous surveillance, along with stricter biosecurity measures, is imperative to limit transmission within poultry and mitigate the risk of zoonotic spillover events.

## Conclusion

G57-like viruses possess evolutionary adaptations which enhance their ability to circulate and persist in terrestrial poultry. These viruses also possess acid-stable HA glycoprotein and mammalian adaption markers, which can increase the chances of their zoonotic spillover to humans. The co-circulation of these viruses with other H9N2 lineages and other influenza subtypes in the Asia Pacific region poses an imminent threat to terrestrial poultry, as it likely generates reassortant viruses with the potential to infect humans.

## Acknowledgements

The authors are thankful for the support provided by the Animal Services team at the Pirbright Institute during transmission studies and to Thomas Peacock, Joe O’Boyle and Toby Carter for their feedback on the manuscript.

---

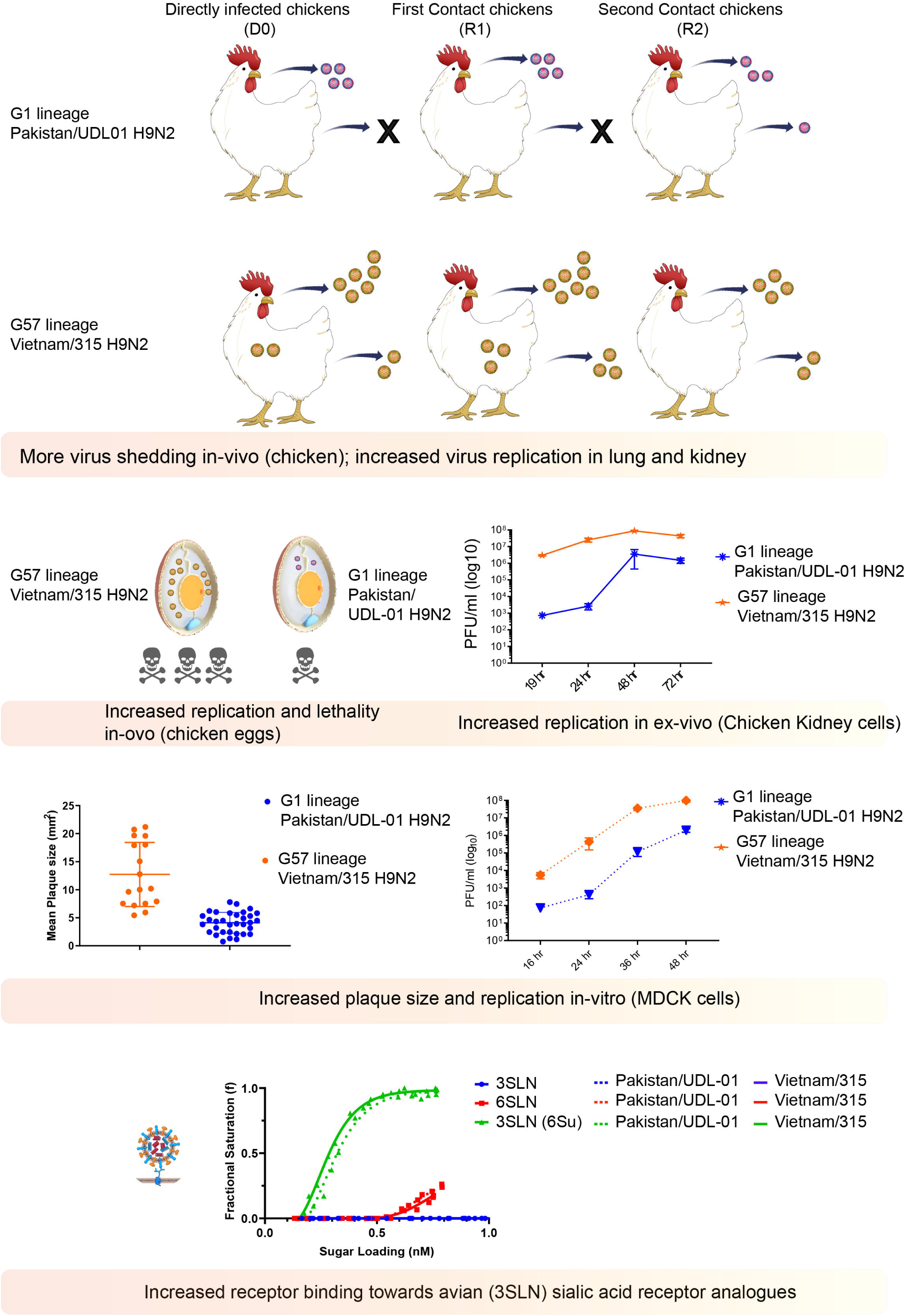

