## Supplementary files for "The G57 genotype of the BJ/94-like H9N2 lineage exhibits increased replication and virulence in chickens compared to the G1 Middle East Group B lineage"

Supplementary data

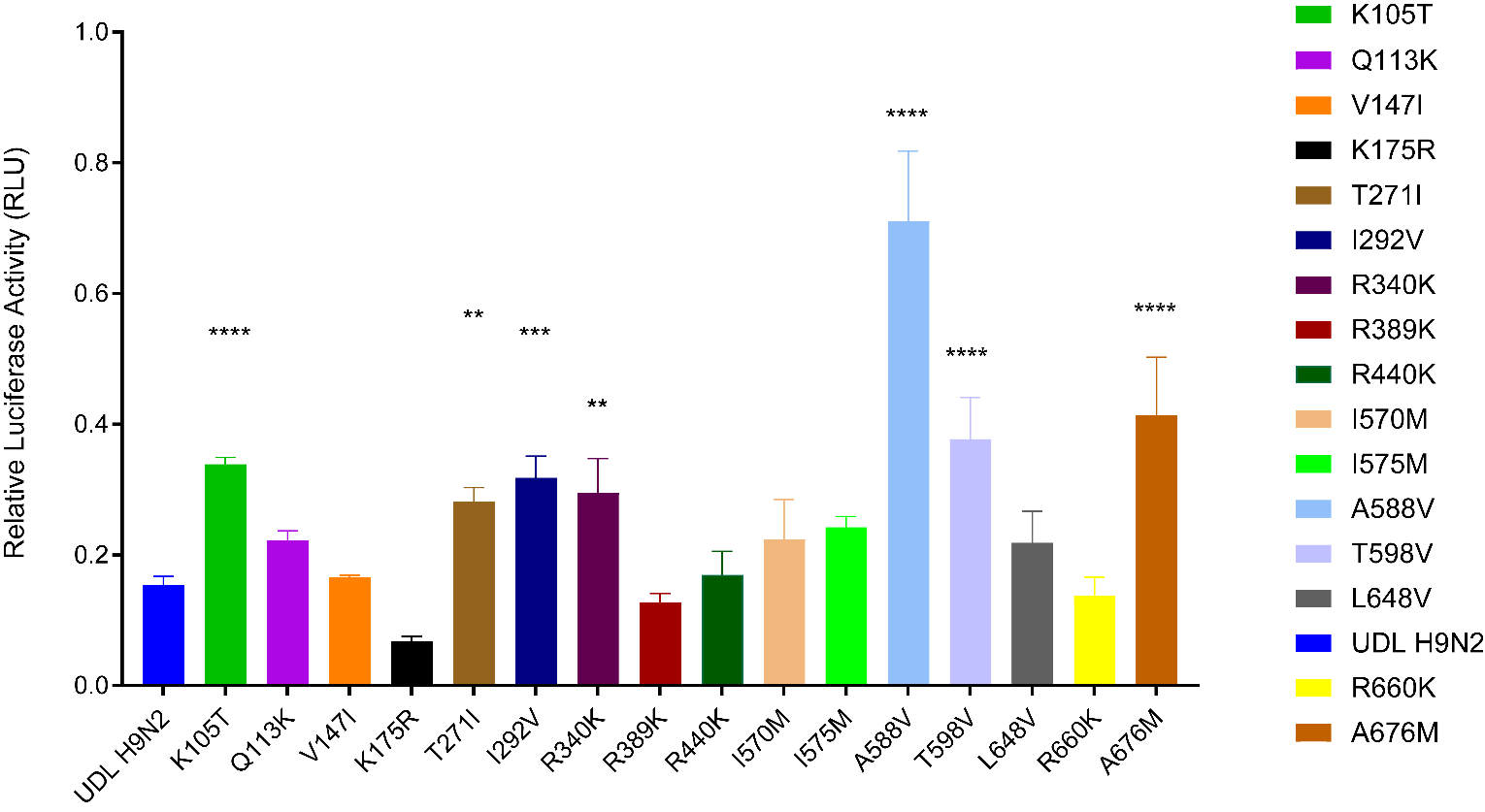

**Supplementary Figure S1: Minireplicon assays of the ribonucleoprotein (RNP) complexes of Pakistan/UDL-01 PB2 mutants.**

The RNP complexes were reconstituted by transfecting human HEK-293T cells and incubating them at 37°C. Luciferase activities were measured 24 h posttransfection. RNP complexes without PB1 served as negative controls (not shown). The data shown are representative of two independent experiments. Ordinary one-way ANOVA was carried out by comparison with Pakistan/UDL-01 H9N2.

** indicates P <0.005; *** indicates P < 0.0005; ****, P < 0.0001.

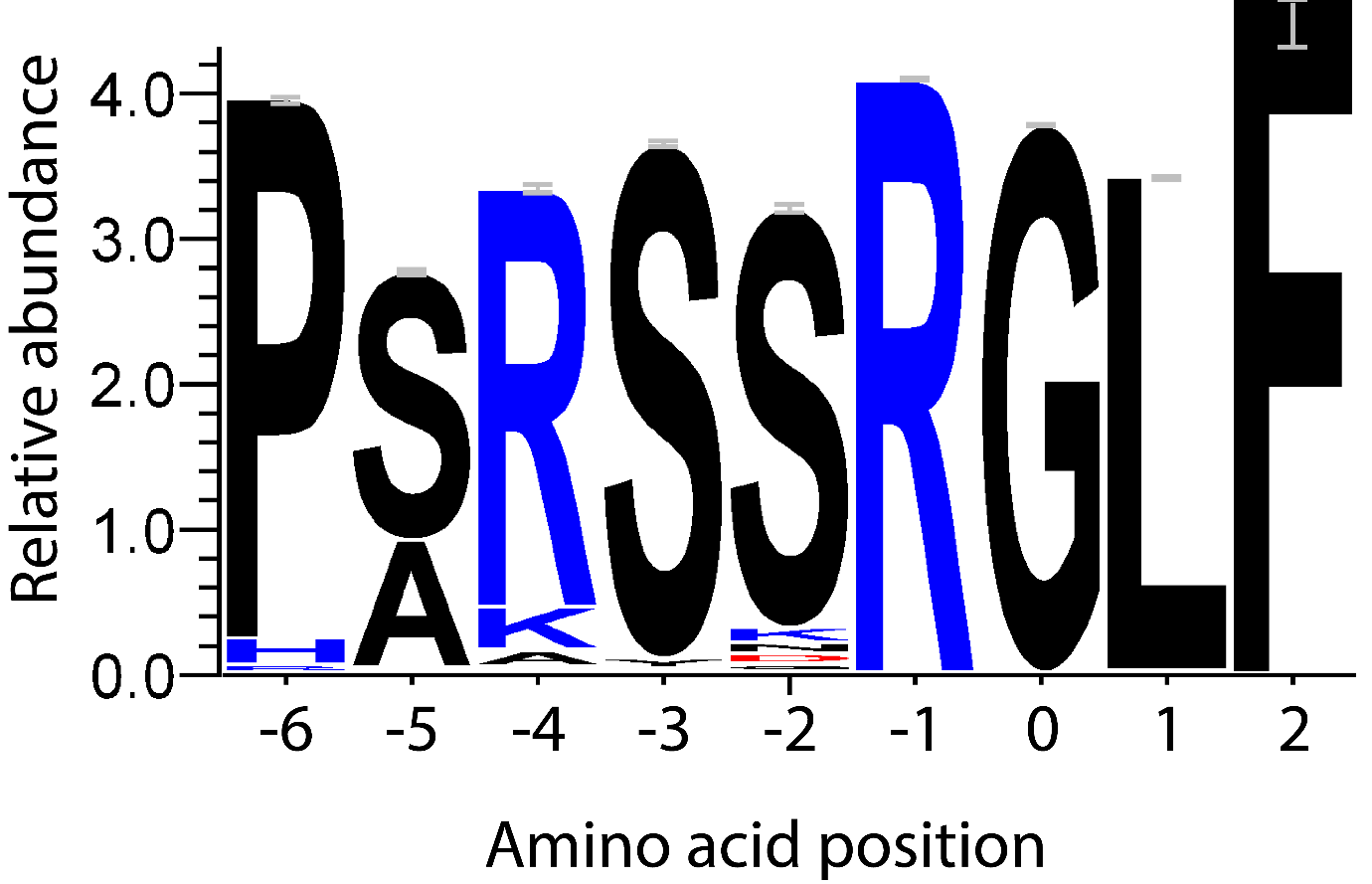

**Supplementary Figure S2: Amino acid analysis of H9HA cleavage site shown as a sequencing logo.** Each logo consists of a symbol or a stack of symbols representing an amino acid at the H9HA cleavage site. The overall height of the stack (shown as bits) indicates the sequence conservation at that position. The height of the symbols within each stack represents the relative frequency of each amino acid at that position. Amino acids are coloured according to their chemical properties. The analysis included 12910 H9HA sequences retrieved from GISAID. The weblogo was created using Weblogo 3 (webversion).

Crooks GE, Hon G, Chandonia JM, Brenner SE WebLogo: A sequence logo generator, Genome Research, 14:1188-1190, (2004)
